# MITE-based drives to transcriptional control of genome host

**DOI:** 10.1101/108332

**Authors:** Luis María Vaschetto

## Abstract

In a recent past, Transposable Elements (TEs) were referred as selfish genetic components only capable of copying themselves with the aim to increase the odds that will be inherited. Nonetheless, TEs have been initially proposed as positive control elements acting in synergy with the host. Nowadays, it is well known that TE movement into genome host comprise an important evolutionary mechanism capable to produce diverse chromosome rearrangements and thus increase the adaptive fitness. According to as insights into TE functioning are increasing day to day, the manipulation of transposition has raised an interesting possibility to setting the host functions, although the lack of appropriate genome engineering tools has unpaved it. Fortunately, the emergence of genome editing technologies based on programmable nucleases, and especially the arrival of a multipurpose RNA-guided Cas9 endonuclease system, has made it possible to reconsider this challenge. For such purpose, a particular type of transposons referred as Miniature Inverted-repeat Transposable Elements (MITEs) has shown a series of interesting characteristics for designing functional drivers. Here, recent insights into MITE elements and versatile RNA-guided CRISPR/Cas9 genome engineering system are given to outline an effective strategy that allows to deploy the TE potential for control of the host transcriptional activity.

## Introduction

Over last decades, diverse genome engineering strategies manipulating the functioning of Transposable Elements (TE) have been intended to achieve site-specific integration of foreign DNA into host genomes. With this aim, the naturally occurring mechanism of transposition was exploited to develop corresponding vectors that allow efficient gene transference and effective integration into genome insertion sites^1–3^. Moreover, the zinc-finger (ZF) and transcription activator-like effector (TALE) programmable nucleases, which can be efficiently redesigned to target specific genome sequences, have been also employed for the development of tools that efficiently achieve genome engineering integration. ZF and TALE-based constructs display important applications in genome engineering, reverse genetics and targeting transgenic integration strategies^4,5^. Lastly, one RNA-guide system based on prokaryotic Type II clustered regularly interspaced short palindromic repeats (CRISPR) and CRISPR-associated Cas9 endonuclease (CRISPR/Cas9) immune system has recently emerged as more efficient, easy to use and low cost genome editing tool able to produce desired genomic modifications^6–8^ (Figure 1).

**Figure 1.**
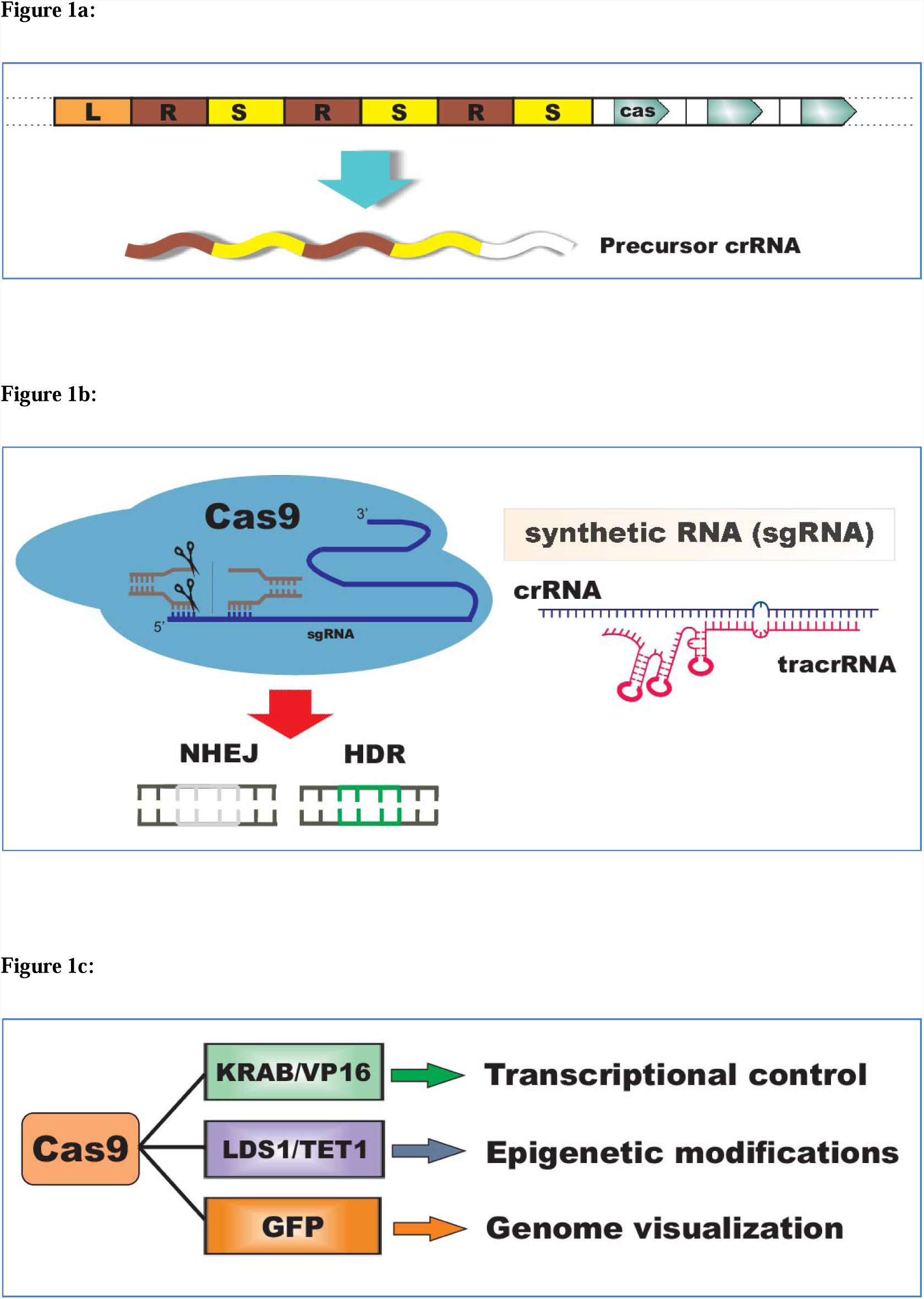
The CRISPR/Cas9 genome editing system. a) The CRISPR locus consist of one leader sequence (L, orange) followed by the CRISPR array containing direct repeats (R, brown) and spacer (Y, yellow) sequences, and CRISPR-associated (Cas) genes. In natural type II CRISPR system, this locus is transcribed as a precursor CRISPR RNA (pre-crRNA) which is processed into mature CRISPR RNAs (crRNAs) molecules that are then bound to transacting CRISPR RNAs (tracrRNAs) in order to direct targeted Cas9-mediated Double Strand Breaks (DSBs) in complementary crRNA sequences. The natural CRISPR system is capable to defend the host against infection of matching crRNA pathogenic sequences b) In CRISPR/Cas9 genome engineering, a synthetic single-guide RNA (sgRNA) that combines the crRNA and tracrRNA molecules is employed to mediate Cas9 DSB cleavage at targeted genomic sites which are subsequently repaired via non-homologous end-joining (NHEJ) or homology-directed (HDR) competing cellular DNA repair pathways. By HDR-mediated repair it is possible to achieve targeted integration of desired template sequences and induce site-specific genome changes. The only requirement of the system is one short Cas9 recognition site known as Protospacer-Adjacent Motif (PAM). c) Alternative Cas9-based chimeric fusion systems can be also designed in order to achieve efficient transcriptional targeting, site-specific epigenetic modifications, cell imagining, etc.

The CRISPR/Cas9-mediated genome editing system use a synthetic single guide RNA (sgRNA) to produce targeted Cas9 DNA Double Strand Breaks (DSBs) that are subsequently rearranged by endogenous DNA repair mechanisms^9,10^. As consequence of this, the DSBs generated are repaired by either both non-homologous end-joining (NHEJ) or homologous recombination (HDR) DNA repair pathways. Depending on repair mechanism activated, site-specific modifications involving gene disruption, gene replacement, or nucleotide substitution may be efficiently generated. NHEJ is prone to produce indel mutations at target sites, while by use of DNA templates harboring desired sequences it is possible generate specific genome modifications by activation of HDR-mediated repair^11^. Alternative genome engineering strategies consisting of a nuclease-dead Cas9 (dCas9) and Cas9 nickases fused with modular transcriptional domains (activators and/or repressors), chromatin-remodelers and fluorophores also enable efficient transcriptional control, site-specific chromatin modifications and visualization of *loci*, respectively^12^. The RNA-guided CRISPR/Cas9 system has dramatically improved our ability for both *in vitro* and *in vitro* genome editing from many organisms, thereby being increasingly employed in biotechnology and therapeutics.

### RNA-guided Cas9 engineering system for design of homing endonuclease gene drives

Diverse endogenous genome elements including, among others, homing endonuclease genes (HEG) and Transposable Elements (TEs) are capable to exploit the host machinery in order to increase the odds that they will be inherited. Such elements were previously referred as “gene drives”, or well parasitic genome players able to spread in the genome host by copying themselves into target sequences^13–15^. Since the beginning, it was believed that these elements might be used to design effective genome engineering strategies that allow to co-opt endogenous host molecular mechanisms. With this aim, Burst^13^ suggested an interesting, multi-talent and promising procedure based on homing endonuclease gene (HEG) drives that would allow to address a wide range of ecological and biotechnological issues (Figure 2); however, several technical constraints hindered its effective design. Even though the Burst’ proposal had not be implemented, his revolutionary idea predicted the development of future outstanding applications. Fortunately, the emergence of the adaptable CRISPR/Cas9 genome editing system has overcome methodological restrictions, thereby Cas9-based drives already successively engineered to bias inheritance in favor of particular yeast (*Saccharomyces cerevisiae*) drives constructs^16^, which exhibited remarkable high transmission rates (99%). Even more importantly, strategies based on CRISPR/Cas9-mediated gene drives has demonstrated have potential to address the evolution of natural populations^17^, whereby displaying the outstanding power that these emerging technologies possess and, besides, underlying an imperative need for public debate before each effective use^18,19^.

**Figure 2.**
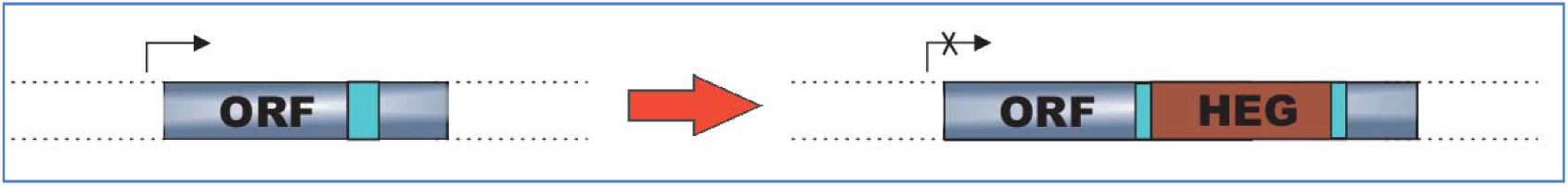
Burst’s proposal (2003) based on targeted homing endonuclease gene (HEG) drives. Genome engineering HEG drives may spread in the genome host via targeted integration into recognition sites (light blue), thereby activating recombination while simultaneously preventing the cleavage of chromosomes that carry them. Following site-specific HEG insertion, it is expected that HEG-based gene drives to be able to modify host cellular functions (in this case, causing loss of function by gene disruption).

### The CRISPR system is associated with the functioning of Transposable Elements

Several parallelisms between the CRISPR/Cas9 system and the eukaryotic RNA interference (RNAi) mechanism have been interestingly discovered^20–22^. In nature, CRISPR *loci* contain arrays of Direct Repeats (DR) associated with spacers sequences which likely derived from bacteriophages or plasmids (see Fig. 1a). The CRISPR DRs act as Cas9-mediated cleavage sites, whereas invasion-acquired spacers mediate immunologic responses to host reinvasion, thereby mimicking eukaryote posttranscriptional gene silencing (PTGS) RNAi mediated by small RNAs (sRNAs). Likewise CRISPR DRs, TEs may also be flanked by cleavage sites recognized by particular enzymes (transposases) that mediate their own transposition. Essentially, there are two types of TE flanking sequences referred as Long Tandem Repeats (LTRs) or Terminal Inverted Repeats (TIRs), which allow their classification into Class I and Class II TEs, respectively. Alternatively, TEs are also categorized according to their transposition intermediates into RNA-mediated (Class I) and DNA-mediated (Class II) elements, which are transposed through “copy-paste” and “cut-paste” mechanisms, respectively (for review see Casacuberta and Santiago^23^). Remarkably, flanking CRISPR DRs derived from insertions of particular Class II TEs known as Miniature Inverted-repeat Transposable Elements (MITEs) have been discovered^24^. MITEs are non-autonomous DNA (Class II) TEs usually composed by ~100-800 pb, flanked by TIRs and adjacent to Tandem Duplication Sites (TDSs) (Figure 3). Resembling to the functioning of CRISPR systems during bacteriophage infection, MITEs also can generate sRNAs that mediate the RNAi silencing pathway during stress responses and environmental (hormone) signals^25^. Altogether, it suggests that both evolution and functioning of natural CRISPR systems are related to the behavior of TEs in relation to their hosts. Indeed, MITE-derived CRISPR DRs indicate that site-specific TE insertions have contributed to evolution of CRISPR arrays^24^. Diverse types of TEs exhibit preference insertional targets, whereby hot spots for many TE families have been correspondingly characterized^26–28^. In the case of MITEs, these elements often exhibit preferential transposition into both AT dinucleotide and ATT trinucleotide genome signatures^29,30^.

**Figure 3.**
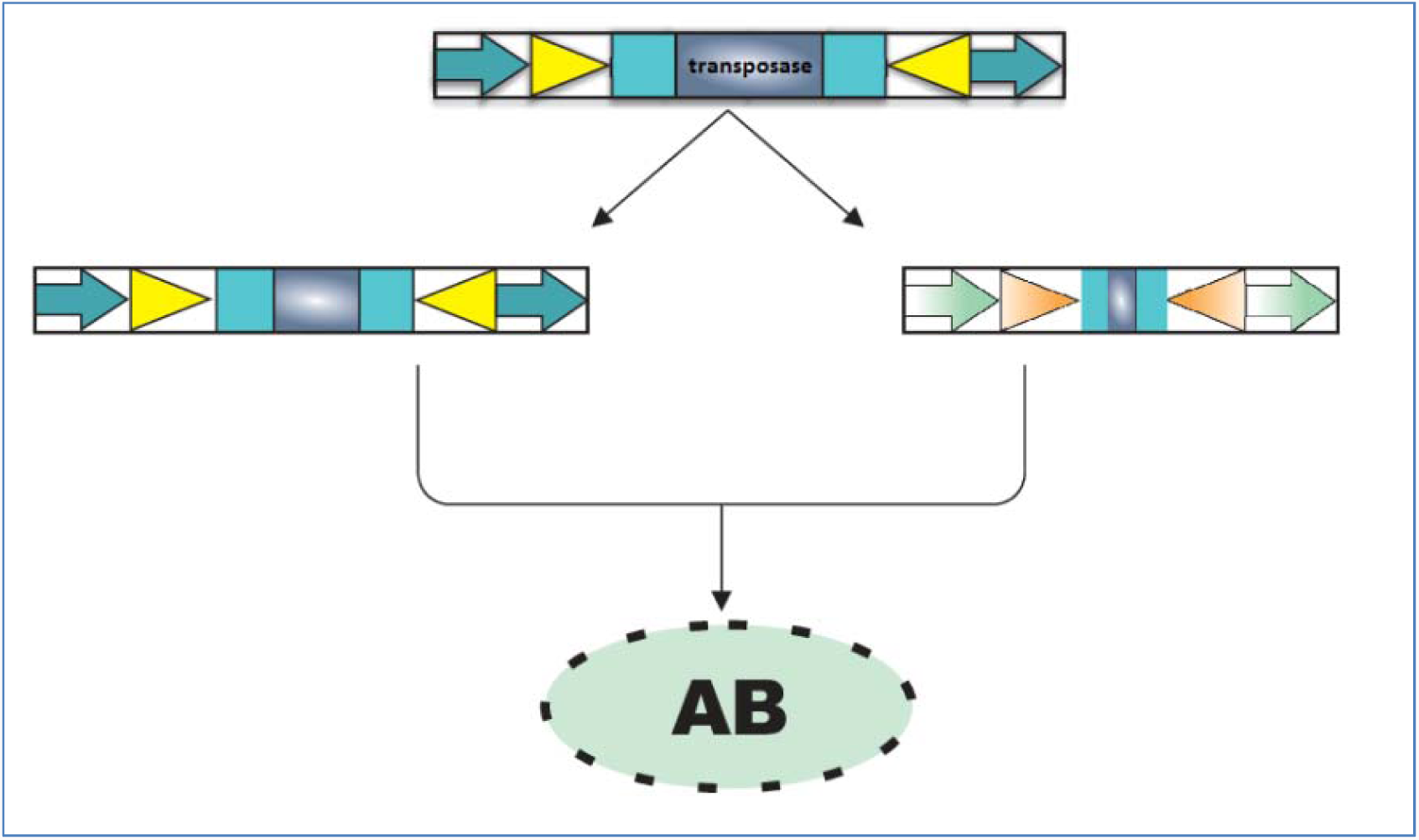
Miniature Inverted-Repeat Transposable Elements (MITEs). MITE elements contain Terminal Inverted Repeats (TIRs, triangles) adjacent to Tandem Duplication Sites (TDSs, arrows). It is believed that MITEs were originated from deletion derivatives of Class II TEs that losing the ability to drive transposition (ie., transposases and functional enzymes) whereas maintaining the transpositional activity. MITE-derived sequences may not contain perfect TIRs or even lost the TDSs. An amplification burst (AB) of MITEs can dramatically increase the number of these elements in one host genome.

From an evolutionary point of view, the transposition mechanism by which TEs are scattered into genome hosts challenged the mistaken concept that considered to genome as one fixed, immutable entity. Nowadays, it is well known that transposition comprise an important adaptive mechanism capable of providing both hereditable and non-hereditable variability, source of critical phenotypic plasticity. Under particular environmental conditions, active transposases mediate transposition and it may positively affect gene expression in the host, thereby generating more adaptable phenotypes. For example, Class I retrotransposons sequences are activated during human neuronal differentiation and consequent amplification triggers chromosome arrangements capable of conferring somatic plasticity^31,32^. In maize, Qüesta and colleagues^33^ suggested that UV-B-radiation induce the transposition via modulation of chromatin structure and thus generate variation in the genome. Moreover, in *Arabidopsis,* it has been showed that stress heat conditions induce the transposition of the ONSEN (Class I) copia-like retrotransposon and TE accumulation was encouraged in plant small intereference RNA (siRNA) deficient mutants^34,35^, whereby evidencing the critical role of RNAi pathways during stress-mediated transposition. Indeed, TE mobilization and epigenetic gene regulation mechanisms are deeply interconnected to each other, and nowadays it is well known that transposition may result in epigenetic modifications associated with more adaptable stress-tolerant phenotypes^36–38^.

### Control of host transcription by induced TE drives

The targeted insertional mutagenesis has emerged as an important strategy for deciphering the gene function by inducing large-scale mutations into genomic loci of interest. In cancer modeling, DNA *Sleeping Beauty* TE-based systems have been used to induce somatic-specific mutagenesis and thus to identify essential genes involved in tumorgenesis^39^. In plants, a transposon-tagging tool for genome-wide analysis based on the rice *mPing* MITEs was used in transgenic soybean^40^, showing this system preferential insertion for both nearby genes and AT-rich sequences. Hancock and colleagues^40^ observed an increased transpositional activity during specific developmental stages (cotyledon vs. globular stage), suggesting interestingly that insights into the developmental regulation pathways involved might be used to control transposition. Both results indicate how the genome fluidity can be efficiently manipulated via TE-based systems. In this sense, new molecular breeding strategies based on the induced activation of Class I retrotransposons have also already been suggested^41^ since, as mentioned before, TEs represent an important source of phenotype plasticity which is particularly interesting in view of the potentiality for addressing transposition through application of external stimulus. As consequence of this we are currently able to engineer TE-based strategies in order to induce transcriptional control of target genes (Figure 4). For such purpose, the versatile CRISPR/Cas9 genome editing system can be used to achieve HDR-mediated preferential insertion upstream of target genes whose transcriptional regulation is desired. Alternatively, RNA-guided genome engineering constructs based on the use of Cas9 fused with transposase enzymes might be likewise employed to direct the insertion of target sequences. Interestingly, targeted transposition by using vector delivery systems consisting of the *piggyBac* transposase fused with ZF/TALE nucleases have been already reported^42–44^. Conclusively, efficient genome editing tools as the multipurpose CRISPR/Cas9 system may be used to address the mechanism of transposition at same time that inducible (environment/stress) conditions allow the simultaneous TE activation, thereby being possible the design of targeted and switchable genome engineering strategies to control the transcription.

**Figure 4.**
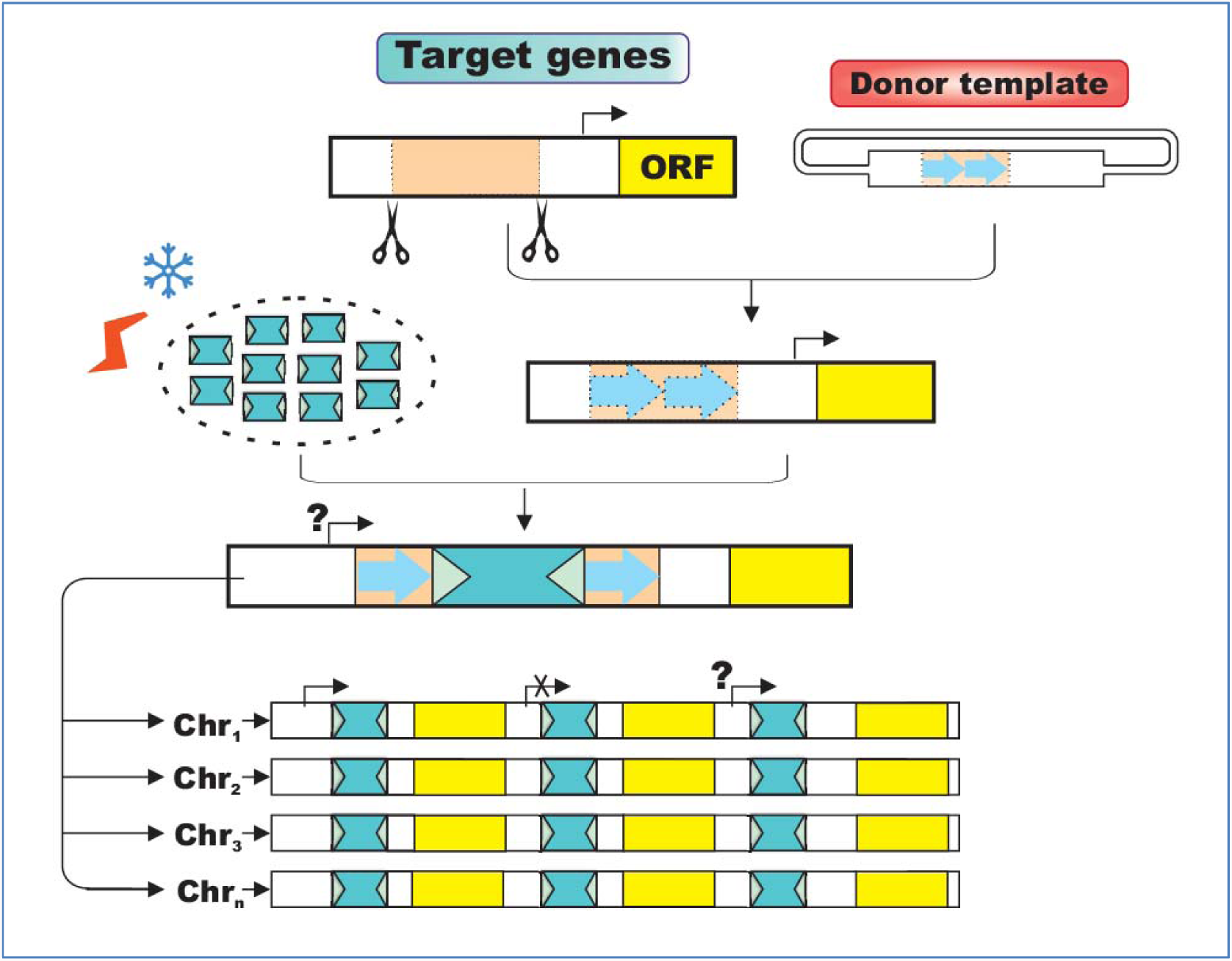
A MITE-based strategy to drive the transcription of target genes. Induced (environmental/stress) conditions would trigger the amplification of MITEs. By taking advantage of the preferential integration at AT/ATT nucleotide sequences, the HDR-mediated CRISPR/Cas9 genome editing technology can be used to generate hot spots into targeted loci which would be occupied by amplified MITEs, thereby targeting integration into host genomes. Such genome engineering strategy would make use of the dual essential property of MITEs to either up- or down-regulate the transcription of nearby genes according to both specific localization of the insertion and regulatory motifs that they may contain.

In order to understand how TE-based strategies could be used for effective control of transcription, insights into the particular type of TE should be mandatory. In this work, MITE elements are described, since these Class II DNA transposons exhibit a series of particular characteristics that make them especially suitable for designing TE-based transcriptional drives. 1- MITEs are abundant repeat elements in eukaryote genomes and they play critical roles during genome evolution^45,46^. For example, MITE elements represent the most usual type of TEs in rice genome^47^. 2- Insertion of MITEs can both upregulate and downregulate the expression of nearby genes^48^. 3- The high number of MITE elements as well as the high level of sequence conservation among MITE subfamilies suggests that they have been amplified from few elements, which is characteristic of Class I TEs^23^. 4- Several active MITEs including, among others, *mPing* and *mGing* in rice^30,49^, *Stowaway* in potato^50^ and *AhMITE1* in peanut^51^ have been discovered. 5- MITE amplification can be efficiently induced under appropriate conditions^49^. 6- MITEs exhibit targeted integration and are preferentially inserted close to genes^45,52^. For instance, the *Tourist-like* and *Stowaway-like* plant MITE superfamilies exhibit preferential insertion into particular TA and TAA target sequences, respectively^47^. 7- Since they do not possess the enzymes required for their own transposition, MITEs are usually shorter compared to Class I and autonomous DNA TEs, thereby being more easily to be manipulated for designing efficient TE-based genome engineer strategies.

In silico analyses show that MITE sequences are involved with RNA-mediated gene regulation^53^, it either by hairpin-like miRNA precursors (pre-miRNA)^54^ or by siRNA biogenesis^55^. In addition, MITE insertions can also mediate gene silencing by the establishment of epigenetic hallmarks such as, for example, repressive DNA and histone methylation. Alternatively, MITE-mediated gene upregulation have been also reported, thereby gene activation through regulatory motifs or epigenetic-dependent mechanisms has been proposed^36,56–59^. In crops, these elements can modulate the transcriptional activity of essential genes^25,38,58,60,61^ and, therefore, MITE insertions may represent an important source of genetic variation to be considered for the improvement of agronomic traits (eg., hydric stress tolerance, yield, disease resistance, quality, etc.). Moreover, genome engineering technologies as, for example, transgenesis, mutagenesis, etc., have been also widely adopted as useful tools for crop improvement. In contrast, MITE-based transcriptional drives might represent an important advantage because this strategy is the only one that would show induced non-hereditable control of transcriptional activity (both up and down), being applicable to different cells, tissues and organs.

## Concluding remarks

Most eukaryotic genomes are littered with TEs, and nowadays it is well known these elements have played critical roles in the evolution of genome host. Although their mode of action resemble to selfish parasitic elements, TE dispersal may also be beneficial for their hosts. Recent insights into the biology of TEs retake an initial position which postulated that environment-induced TE dispersion might comprise an important adaptive mechanism^62^, and nowadays it is recognized as a regulatory mechanism for both allocating gene motifs and genome sequences involved in the establishment of epigenetic profiles. Indeed, the amplification of TEs induced under certain conditions may offer significant benefits to the host by modulating expression of non-linked genes involved in the same gene regulatory pathways, even also having the potential to generate de novo regulatory networks^63^. The currently active *mPing* MITEs represent a clear example of how TEs can act synergistically with the host to modulate its functioning. The *mPing* family is present in high copies in the rice genome and exhibit preferential insertion into AT-gene-rich regions, avoiding exons while simultaneously choosing promoter regions. These elements are able to both upregulate or downregulate the expression of genes according to the localization of the insertion, although more than 80% of *mPing* insertions did not exhibit detectable effects on expression of nearby genes^64^. Interestingly, Naito and colleagues^65^ proposed that active *mPing* elements should be benign to hosts, whereby consequent amplification might result in the selective control of gene expression. On the other hand, since it has been shown that transgenic *mPing*-based transposon tagging systems can also remain active in the soybean genome^40^, the functional characterization of *mPing* MITEs would allow design effective genome engineering strategies involving not only endogenous but also exogenous control of the host functions.

Transposon-based vectors are very useful and effective tools for achieving targeted mutagenesis (gene disruption/replacement). Moreover, the emergence of the versatile CRISPR/Cas9 genome editing system has enabled to overcome long-standing technical limitations in order to give the next step towards the design of more efficient TE-based targeting systems. The preferential integration induced through application of particular stress/environmental treatments represent a key milestone to maximize the versatility of such systems, since it allows to exploit at will the natural potential that TEs have to control the host. Further advances on CRISPR research will allow to develop new and original approaches that surely overcome the challenges associated to design besides those that RNA-guided systems themselves pose (off-target effects, targeting specificity, etc). As has been exposed in this article, MITEs exhibit a series of well-defined characteristics worthy to be considered the engineering design of environment-induced TE-based drives. In particular, it will depend on the ability to induce effective activation, whereby *mPing* elements have been found to be rough diamonds, since their amplification can be efficiently triggered by application of different treatments such as cell culture^30,66^, hybridization^67^ or pressurization^68^. In resemblance with HEG-mediated gene drive strategies, issues related to cutting specificity, copying efficiency and stability of TE drives should be also considered, being certainly the CRISPR/Cas9-based genome engineering tools proper to address all these methodological challenges. In the case to achieve cell-specific regulation of gene expression, TE-based drives will represent an efficient and non-hereditable method of transcriptional control.

It should be noted that transcriptional drives would allow to address important ecological and biotechnological issues. In crops, the climatic fluctuations associated with the global warming require the design of molecular breeding strategies tailored to optimize plant development under such scenarios of change^37^. In this regard, it has been already proposed that the TE amplification may represent a solution for generating functional genetic diversity in the face of ever-changing environments^64^. Thus, induced activation of TE-based drives represent an innovative approach to control the transcriptional activity of target genes whose expression has been positively correlated with the development of quantitative (complex) traits such as biomass, seed composition, yield, disease resistance, etc. Moreover, it would be feasible to engineer in crops ecological TE drives which activated under excessive precipitation conditions upregulate the activity of target genes influencing evapotranspiration efficiency, thereby designing a biologically versatile and useful strategy to drain the surplus of water within floodplain areas. On the other hand, since it has been found that TEs are associated with diverse genome host mechanisms including, among others, telomere maintenance^69^, chromosomal rearrangements^70^, gene duplication^71^ and epigenetic regulation^72^, similar strategies could be eventually applied in order to exploit such functions. At root the impulse to develop these outstanding applications it is certainly likely that, likewise MITE-mediated transcriptional drives, genome engineering models based on other types of TEs will be similarly proposed as to the extent that our knowledge on their functioning gain success to address it. In conclusion, even though TE-based drives pose challenges to overcome, the implementation of effective strategies that exploit such systems should be a short-term issue.

## References

1. Szabo, M. et al. Transposition and targeting of the prokaryotic mobile element IS30 in zebrafish. FEBS Lett. 550, 46–50 (2003)

2. Yant, S. R., Huang, Y., Akache, B. & Kay, M. A. Site-directed transposon integration in human cells. Nucleic acids res 35, e50. (2007).

3. Palazzoli, F., Testu, F. X., Merly, F. & Bigot, Y. Transposon tools: worldwide landscape of intellectual property and technological developments. Genetica 138, 285–299 (2010).

4. Walsh, R. M. & Hochedlinger, K. A variant CRISPR-Cas9 system adds versatility to genome engineering. P Natl Acad Sci USA 110, 15514–15515 (2013).

5. Bortesi, L., & Fischer, R. The CRISPR/Cas9 system for plant genome editing and beyond. Biotechnol adv 33, 41–52 (2015).

6. Cong, L. et al. Multiplex genome engineering using CRISPR/Cas systems. Science 339, 819–823 (2013).

7. Ran, F. A. et al. Double nicking by RNA-guided CRISPR Cas9 for enhanced genome editing specificity. Cell 154, 1380–1389 (2013).

8. Cox, D. B. T., Platt, R. J. & Zhang, F. Therapeutic genome editing: prospects and challenges. Nat med 21, 121–131 (2015).

9. Jinek, M. et al. A programmable dual-RNA-guided DNA endonuclease in adaptive bacterial immunity. Science 337, 816–821 (2012).

10. Doudna, J. A. & Charpentier, E. The new frontier of genome engineering with CRISPR-Cas9. Science 346, 1258096 (2014).

11. Sakuma, T. & Yamamoto, T. CRISPR/Cas9: the leading edge of genome editing technology. In Targeted Genome Editing Using Site-Specific Nucleases (pp. 25–41). Springer Japan (2015).

12. Li, X., Wu, R. & Ventura, A. The present and future of genome editing in cancer research. Hum Genet 135, 1083–1092 (2016).

13. Burt, A. Site-specific selfish genes as tools for the control and genetic engineering of natural populations. Proc. Biol. Sci. 270, 921–928 (2003).

14. Rasgon, J. L. & Gould, F. Transposable element insertion location bias and the dynamics of gene drive in mosquito populations. Insect mol biol 14, 493–500 (2005).

15. Adelman, Z. N. & Tu, Z. Control of mosquito-borne infectious diseases: sex and gene drive. Trends parasitol, 32, 219–229 (2016).

16. DiCarlo, J. E. et al. RNA-guided gene drives can efficiently bias inheritance in wild yeast. bioRxiv doi:http://dx.doi.org/10.1101/013896 (2015).

17. Hammond, A. et al. A CRISPR-Cas9 gene drive system targeting female reproduction in the malaria mosquito vector Anopheles gambiae. Nat biotechnol 34, 78–83 (2016).

18. Esvelt, K. M., Smidler, A. L., Catteruccia, F. & Church, G. M. Concerning RNA-guided gene drives for the alteration of wild populations. Elife 3, e03401 (2014).

19. Akbari, O. S. et al. Safeguarding gene drive experiments in the laboratory. Science 349, 927–929 (2015).

20. Mojica, F. J. M. et al. Intervening sequences of regularly spaced prokaryotic repeats derive from foreign genetic elements. J Mol Evol 60, 174–182 (2005).

21. Bolotin, A. et al. Clustered regularly interspaced short palindrome repeats (CRISPRs) have spacers of extrachromosomal origin. Microbiology, 151, 2551–2561 (2005).

22. Barrangou, R. et al. Advances in CRISPR-Cas9 genome engineering: lessons learned from RNA interference. Nucleic acids res 43, 3407–19 (2015).

23. Casacuberta, J. M. & Santiago, N. Plant LTR-retrotransposons and MITEs: control of transposition and impact on the evolution of plant genes and genomes. Gene 311, 1–11(2003).

24. Mai, G., Ge, R., Sun, G., Meng, Q. & Zhou, F. A comprehensive curation shows the dynamic evolutionary patterns of prokaryotic CRISPRs. Biomed res int. doi: 10.1155/2016/7237053 (2016).

25. Yan, Y. et al. Small RNAs from MITE-derived stem-loop precursors regulate abscisic acid signaling and abiotic stress responses in rice. Plant J 65, 820–828 (2011).

26. Castilho, B. A. & Casadaban, M. J. Specificity of mini-Mu bacteriophage insertions in a small plasmid. J bacteriol 173, 1339–1343 (1991).

27. Mandal, P. K. et al. Identification of insertion hot spots for non-LTR retrotransposons: computational and biochemical application to Entamoeba histolytica. Nucleic acids res 34, 5752–5763 (2006).

28. Feng, G., Leem, Y. E. & Levin, H. L. Transposon integration enhances expression of stress response genes. Nucleic acids res 41, 775–89 (2013).

29. Jiang, N. & Wessler, S. R. Insertion preference of maize and rice miniature inverted repeat transposable elements as revealed by the analysis of nested elements. Plant Cell, 13, 2553–2564 (2001).

30. Jiang, N. et al. (2003) An active DNA transposon family in rice. Nature 421, 163–167.

31. Singer, T. et al. LINE-1 retrotransposons: mediators of somatic variation in neuronal genomes?. Trends neurosci 33, 345–354 (2010).

32. Baillie, J. K. et al. Somatic retrotransposition alters the genetic landscape of the human brain. Nature 479, 534–537 (2011).

33. Qüesta, J. I., Walbot, V. & Casati, P. Mutator transposon activation after UV-B involves chromatin remodeling. Epigenetics 5, 352–363(2010).

34. Ito, H. et al. An siRNA pathway prevents transgenerational retrotransposition in plants subjected to stress. Nature 472, 115–119 (2011).

35. Matsunaga, W., Kobayashi, A., Kato, A. & Ito, H. The effects of heat induction and the siRNA biogenesis pathway on the transgenerational transposition of ONSEN, a copia-like retrotransposon in Arabidopsis thaliana. Plant Cell Physiol 53, 824–833 (2012).

36. Yang, G. et al. A two-edged role for the transposable element Kiddo in the rice ubiquitin2 promoter. Plant Cell 17, 1559–1568 (2005).

37. Hou, J., Long, Y., Raman, H., Zou, X., Wang, J., Dai, S., … & Meng, J. (2012). A Tourist-like MITE insertion in the upstream region of the BnFLC. A10 gene is associated with vernalization requirement in rapeseed (Brassica napus L.). BMC plant biology, 12(1), 1.

38. Castelletti, S., Tuberosa, R., Pindo, M. & Salvi, S. A MITE transposon insertion is associated with differential methylation at the maize flowering time QTL Vgt1. G3 4, 805–812 (2014).

39. Molyneux, S. D. et al. Human somatic cell mutagenesis creates genetically tractable sarcomas. Nat genet 46, 964–972 (2014).

40. Hancock, C. N. et al. The rice miniature inverted repeat transposable element mPing is an effective insertional mutagen in soybean. Plant physiol 157, 552–562. (2011).

41. Paszkowski, J. Controlled activation of retrotransposition for plant breeding. Curr opin biotech 32, 200–206 (2015).

42. Kettlun, C. et al. Manipulating piggyBac transposon chromosomal integration site selection in human cells. Mol Ther 19, 1636–1644 (2011).

43. Li, X. et al. piggyBac transposase tools for genome engineering. Proc Natl Acad Sci USA 110, E2279–E2287 (2013).

44. Owens, J. B. et al. Transcription activator like effector (TALE)-directed piggyBac transposition in human cells. Nucleic Acids Res 41, 9197–9207 (2013).

45. Lu, C. et al. Miniature inverted-repeat transposable elements (MITEs) have been accumulated through amplification bursts and play important roles in gene expression and species diversity in Oryza sativa. Mol biol evol 29, 1005–1017 (2012).

46. Ye, C., Ji, G. & Liang, C. detectMITE: A novel approach to detect miniature inverted repeat transposable elements in genomes. Sci rep 6 (2016).

47. Jiang, N., Feschotte, C., Zhang, X. & Wessler, S. R. Using rice to understand the origin and amplification of miniature inverted repeat transposable elements (MITEs). Curr opin plant biol 7, 115–119 (2004).

48. Chen, J., Hu, Q., Lu, C. & Kuang, H. Evolutionary Genomics of Miniature Inverted-Repeat Transposable Elements (MITEs) in Plants. In Evolutionary Biology: Genome Evolution, Speciation, Coevolution and Origin of Life (pp. 157–168). Springer International Publishing (2014).

49. Dong, H. T. et al. A Gaijin-like miniature inverted repeat transposable element is mobilized in rice during cell differentiation. BMC Genomics 13:135 (2012).

50. Momose, M., Abe, Y. & Ozeki, Y. Miniature inverted-repeat transposable elements of Stowaway are active in potato. Genetics 186(1):59–66. (2010)

51. Shirasawa, K. et al. Characterization of active miniature inverted-repeat transposable elements in the peanut genome. Theor Appl Genet 124, 1429–1438(2012).

52. Wessler, S.R., Bureau, T.E. & White, S.E. LTR-retrotransposons and MITEs: important players in the evolution of plant genomes. Curr. Opin. Genet. Dev. 5, 814–821 (1995).

53. Piriyapongsa, J. & Jordan, I. K. Dual coding of siRNAs and miRNAs by plant transposable elements. Rna 14, 814–821(2008).

54. Lorenzetti, A. P., de Antonio, G. Y., Paschoal, A. R. & Domingues, D. S. PlanTE-MIR DB: a database for transposable element-related microRNAs in plant genomes. Funct integr genomic 16, 235–242 (2016).

55. Kuang, H. et al. Identification of miniature inverted-repeat transposable elements (MITEs) and biogenesis of their siRNAs in the Solanaceae: new functional implications for MITEs. Genome res 19, 42–56 (2009).

56. Zhang, Q., Arbuckle, J. & Wessler, S.R. Recent, extensive, and preferential insertion of members of the miniature inverted-repeat transposable element family Heartbreaker into genic regions of maize. Proc. Natl. Acad. Sci. USA 97 1160–1165 (2000).

57. Sun, X. et al. PdMLE1, a specific and active transposon acts as a promoter and confers Penicillium digitatum with DMI resistance. Env microbiol repor 5, 135–142(2013).

58. Li, J., Wang, Z., Peng, H. & Liu, Z. A MITE insertion into the 3′-UTR regulates the transcription of TaHSP16. 9 in common wheat. Crop J 2, 381–387. (2014).

59. Vaschetto, L. M. Miniature Inverted-repeat Transposable Elements (MITEs) and their effects on the regulation of major genes in cereal grass genomes. Mol Breed 36, 1–4 (2016).

60. Patel, M. et al. High-oleate peanut mutants result from a MITE insertion into the FAD2 gene. Theor Appl Genet 108, 1492–1502 (2004).

61. Mao, H. et al. A transposable element in a NAC gene is associated with drought tolerance in maize seedlings. Nat commun 6 (2015).

62. McClintock, B. The significance of responses of the genome to challenge. Science 226, 792–801 (1984).

63. Feschotte, C. Transposable elements and the evolution of regulatory networks. Nat Rev Genet 9, 397–405 (2008).

64. Naito, K. et al. Unexpected consequences of a sudden and massive transposon amplification on rice gene expression. Nature 461, 1130–1134 (2009).

65. Naito, K. et al. mPing: The bursting transposon. Breeding sci 64, 109–114 (2014).

66. Kikuchi, K. et al. The plant MITE mPing is mobilized in anther culture. Nature 421, 167–170(2003).

67. Shan, X. et al. Mobilization of the active MITE transposons mPing and Pong in rice by introgression from wild rice (Zizania latifolia Griseb.). Mol Biol Evol 22, 976–990 (2005).

68. Lin, X. et al. In planta mobilization of mPing and its putative autonomous element Pong in rice by hydrostatic pressurization. J Exp Bot 57, 2313–2323 (2006).

69. Aschacher, T. et al. LINE-1 induces hTERT and ensures telomere maintenance in tumour cell lines. Oncogene 35, 94–104 (2016).

70. Feschotte, C., & Pritham, E. J. DNA transposons and the evolution of eukaryotic genomes. Annu Rev Genet 41, 331–368 (2007).

71. Flagel, L. E., & Wendel, J. F. Gene duplication and evolutionary novelty in plants. New Phytol 183, 557–564 (2009).

72. Slotkin, R. K., & Martienssen, R. Transposable elements and the epigenetic regulation of the genome. Nat Rev Genet, 8, 272–285 (2007).

